# Bundle sheath extensions affect leaf structural and physiological plasticity in response to irradiance

**DOI:** 10.1101/208850

**Authors:** Maria Antonia M. Barbosa, Daniel H. Chitwood, Aristéa A. Azevedo, Wagner L. Araújo, Dimas M. Ribeiro, Lázaro E. P. Peres, Samuel C. V. Martins, Agustin Zsögön

## Abstract

Coordination between structural and physiological traits is key to plants’ responses to environmental fluctuations. In heterobaric leaves, bundle sheath extensions (BSEs) increase photosynthetic performance (light-saturated rates of photosynthesis, *A*_max_) and water transport capacity (leaf hydraulic conductance, *K*_leaf_). However, it is not clear how BSEs affect these and other leaf developmental and physiological parameters in response to environmental conditions. The *obscuravenosa* (*obv*) mutation, found in many commercial tomato varieties, leads to absence of BSEs. We examined structural and physiological traits of tomato heterobaric and homobaric (*obv*) near-isogenic lines (NILs) grown at two different irradiance levels. *K*_leaf_, minor vein density and stomatal pore area index decreased with shading in heterobaric but not in homobaric leaves, which show similarly lower values in both conditions. Homobaric plants, on the other hand, showed increased *A*_max_, leaf intercellular air spaces and mesophyll surface area exposed to intercellular airspace (*S*_mes_) in comparison with heterobaric plants when both were grown in the shade. BSEs further affected carbon isotope discrimination, a proxy for long-term water-use efficiency. BSEs confer plasticity in traits related to leaf structure and function in response to irradiance levels and might act as a hub integrating leaf structure, photosynthetic function and water supply and demand.

**Summary statement:** The presence of bundle sheath extension (BSEs) defines leaves as heterobaric, as opposed to homobaric leaves that lack them. Multiple functions have been proposed for BSEs, but their impact on different environmental conditions is still unclear. Here, we compared a tomato (*Solanum lycopersicum*) homobaric mutant lacking BSEs with its corresponding heterobaric wild-type, grown under two irradiance conditions. We show that the presence of BSEs differentially alters various physiological and anatomical parameters in response to growth irradiance. We propose that BSEs could act as hubs coordinating leaf plasticity in response to environmental factors.

**Article type:** Research article

## Introduction

Leaves are the evolutionary solution to maximize light capture and optimize CO_2_ and water vapour exchange with the atmosphere in land plants. Leaf biochemistry and structure are, therefore, strongly coordinated with photosynthetic performance and hydraulic function. Whereas such coordination is of paramount importance for plant growth and ecological distribution (Nicotra *et al.* 2008; Nicotra *et al.* 2011), it also requires a degree of developmental plasticity to cope with environmental variation given the sessile nature of plants (Schlichting 1986; Valladares *et al.* 2007). The light environment can be highly variable and dynamic, being particularly effective at influencing leaf structure and function (Terashima *et al.* 2001; Terashima *et al.* 2006). Leaf anatomy, in turn, can influence CO_2_ and H_2_O exchanges with the atmosphere (Evans & Poorter 2001; Scoffoni *et al.* 2015). Optimality theory predicts that, under a given set of conditions, all parameters will tend to converge to maximize photosynthesis with the available resources, mainly light, nitrogen and water (Niinemets 2012 and references therein).

Rubisco activity, capacity for ribulose-1,5-bisphosphate regeneration and triose-phosphate export from chloroplasts are key biochemical determinants of net photosynthesis rate (*A*). Photosynthetic carbon assimilation, however, depends not only on the biochemistry of the leaf, but also on its diffusive properties which are strongly dependent on anatomy and morphology (Terashima, Hanba, Tholen & Niinemets 2011; Nunes-Nesi *et al.* 2016). Strong correlations with *A* have been found for stomatal distribution between the adaxial and abaxial faces (*i.e.* amphistomatous or hypostomatous leaves), blade thickness, leaf mass per area, the palisade-to-spongy mesophyll ratio, and the area of mesophyll and chloroplast surfaces facing the intercellular air spaces (Niinemets & Sack 2006). All of these parameters are highly plastic in response to light (Oguchi *et al.* 2003; Oguchi *et al.* 2005; Terashima *et al.* 2011) and potentially affect how water transport and evaporation occur in the leaf (Sack *et al.* 2003; Sack & Frole 2006). The efficiency of water transport through the leaf is measured as *K*_leaf_ (leaf hydraulic conductance) (Sack & Holbrook 2006), which has been shown highly dynamic and able to vary rapidly with time of day, irradiance, temperature, and water availability (Prado & Maurel 2013). Leaf structural traits such as blade thickness, stomatal pore area, lamina margin dissection, among others, have been shown to influence *K*_leaf_ (Sack & Holbrook 2006).

In particular, vein structure and patterning play a critical role in determining both carbon assimilation rate (McAdam *et al.* 2017) and water distribution within plants (Sack *et al.* 2012). Water flow through the leaf occurs via xylem conduits within the vascular bundles, which upon entering the lamina from the petiole, rearrange into major and minor veins. Upon leaving the xylem, water has to transit through the bundle sheath, a layer of compactly arranged parenchymatic cells surrounding the vasculature (Trifiló, Raimondo, Savi, Lo Gullo & Nardini 2016; Scoffoni *et al.* 2017). Bundle sheaths could behave as flux sensors or ‘control centers’ of leaf water transport, and they are most likely responsible for the high dependence of *K*_leaf_ on temperature and irradiance (Leegood 2008; Ohtsuka *et al.* 2018). Vertical layers of colorless cells connecting the vascular bundle to the epidermis are present in many eudicotyledons (Esau 1977). These so-called bundle sheath extensions (BSEs) are most commonly found in minor veins, but can occur in veins of any order depending on the species (Wylie 1943; Wylie 1952). A topological consequence of the presence of BSEs is the formation of compartments in the lamina, which restricts lateral gas flow and thus allows compartments to maintain gas exchange rates independent of one another (Pieruschka, Schurr, Jensen, Wolff & Jahnke 2006; Morison, Lawson & Cornic 2007; Buckley, Sack & Gilbert 2011). Such leaves, and by extension the species possessing them, are therefore called ‘heterobaric’, as opposed to ‘homobaric’ species lacking BSEs (Neger 1918).

Large taxonomic surveys have demonstrated that heterobaric species tend to occur more frequently in sunny and dry sites or in the upper stories of climax forests (Kenzo *et al.* 2007), so it was hypothesized that BSEs could fulfill an ecological role by affecting mechanical and physiological parameters in the leaf (Terashima 1992). Despite some proposed functions for BSEs (mechanical support, increased damage resistance, among others) remain hypothetical (Lawson & Morison 2006; Read & Stokes 2006), other functions have been proven through meticulous experimental work, suggesting that the existence of BSEs could be adaptive (Buckley *et al.* 2011). For instance, lateral propagation of ice in the lamina was precluded by the sclerenchymatic BSEs in *Cinnamomum canphora* L, although this effect has only hitherto been described in this species and could depend on the type and amount of BSEs in the leaf blade (Hacker & Neuner 2007). Hydraulic integration of the lamina was increased by BSEs, which connect the vascular bundle to the epidermis and, therefore, reduce the resistance in the water path between the supply structures (veins) and the water vapor outlets (stomata) (Zwieniecki *et al.* 2007). Lastly, *A* was increased in leaves with BSEs, due to their optimization of light transmission within the leaf blade (Karabourniotis *et al.* 2000; Nikolopoulos *et al.* 2002).

We have previously characterized a homobaric mutant that lacks BSEs in the otherwise heterobaric species tomato (*Solanum lycopersicum* L.) (Zsögön *et al.* 2015). The homobaric mutant *obscuravenosa* (*obv*) reduces *K*_leaf_ and stomatal conductance but does not impact *A*_max_, nor global carbon economy of the plant. Here, we extend our observations to plants grown under two contrasting irradiance levels, which are known to influence leaf structure (Oguchi *et al.* 2003; Oguchi *et al.* 2005; Oguchi *et al.* 2006), *A*_max_ (Evans & Poorter 2001; Shipley 2002) and *K*_leaf_ (Scoffoni *et al.* 2008; Guyot *et al.* 2012; Scoffoni *et al.* 2015). We investigated whether the presence of BSEs could have an impact on the highly plastic nature of leaf development and function in response to different irradiance levels. We hypothesized that homobaric leaves, lacking a key physical feature that increases carbon assimilation and leaf hydraulic integration, would exhibit less plasticity in their response to environmental conditions than heterobaric leaves. By assessing a series of leaf structural and physiological parameters in tomato cultivar Micro-Tom (MT) and the near-isogenic *obv* mutant, we provide evidence of the potential role of BSEs in the coordination of leaf structure and hydraulics in response to growth irradiance. Finally, we analysed whether dry mass accumulation and tomato fruit yield are affected by the presence of BSEs and irradiance in two different tomato genetic backgrounds (cultivars MT and M82). We discuss the potential role of BSEs in the coordination of leaf structure and function in response to the light environment.

## Materials and Methods

### Plant material and experimental setup

Seeds of the tomato (*Solanum lycopersicum* L.) cv Micro-Tom (MT) and cv M82 were donated by Dr Avram Levy (Weizmann Institute of Science, Israel) and the Tomato Genetics Resource Center (TGRC, Davis, University of California, CA, USA), respectively. The introgression of the *obscuravenosa* (*obv*) into the MT genetic background to generate a near-isogenic line (NIL) was described previously (Carvalho *et al.* 2011). The model tomato M82 cultivar harbors the *obv* mutation, so the experiments were performed on F1 lines obtained by crosses between MT and M82. Both F1 lines have 50% MT and 50% M82 genome complement, differing only in the presence or absence of BSEs (described in Table 1). Data were obtained from two independent assays, similar results were found both times. Plants were grown in a greenhouse in Viçosa (642 m asl, 20° 45’ S; 42° 51’ W), Minas Gerais, Brazil, under semi-controlled conditions. Micro-Tom (MT) background plants were grown during the months of May to August of 2016 in temperature of 24/20°C, 13/11h (day/night) photoperiod. Plants in the M82 background were cultivated during the months of September to December of 2016 with temperature of 26/22°C, 12/12h (day/night) photoperiod. Plant cultivation was carried out as described previously (Silva *et al.* 2018). The experiments were conducted in completely randomized experimental design, in 2×2 factorial, consisting of two genotypes, and two irradiance levels (sun and shade). Plants in the ‘sun’ treatment were exposed to greenhouse conditions, with midday irradiance of ~900 μmol photons m^−2^ s^−1^. For the ‘shade’ treatment plants were maintained on a separate bench covered with neutral shade cloth, with a retention capacity of 70% of sunlight (250-300 μmol photons m^−2^ s^−1^).

**Table 1.** Description of the plant material used in this study. Micro-Tom (MT) and M82 are two tomato cultivars that differ in growth habit due mostly to the presence of a mutant allele of the *DWARF* gene, which codes for a key enzyme of the brassinosteroid biosynthesis pathway. The molecular identity of *OBSCURAVENOSA* (*OBV*) is unknown. MT harbors a functional, dominant allele of *OBV*, whereas M82 is a mutant (*obv*). F1 plants are hybrids with a 50/50 MT/M82 genomic complement, differing only in the presence or absence of BSEs. The F1 plants are otherwise phenotypically indistinguishable from the M82 parent.

| Parental genotype | MT | MT-*obv* | M82 | F1 MT×M82 | F1 MT-*obv*×M82 |
| --- | --- | --- | --- | --- | --- |
| *Plant height* |  |  |  |  |  |
| Genotype | *dwarf/dwarf* | *dwarf/dwarf* | *DWARF/DWARF* | *DWARF/dwarf* | *DWARF/dwarf* |
| Phenotype | Dwarf plant | Dwarf plant | Tall plant | Tall plant | Tall plant |
| *BSEs* |  |  |  |  |  |
| Genotype | *OBV/OBV* | *obv/obv* | *obv/obv* | *OBV/obv* | *obv/obv* |
| Phenotype | BSEs (clear veins) | No BSEs (dark veins) | No BSEs (dark veins) | BSEs (clear veins) | No BSEs (dark veins) |

### Plant morphology determinations

Morphological characterization was performed in MT plants 50 days after germination as described (Vicente *et al.* 2015). Specific leaf area (SLA) was calculated through the relationship between leaf area (LA) and dry mass (LDW), as described by the equation SLA (cm^2^ g^−1^) = LA/LDW.

Leaflet outline shape was analysed as described in Chitwood *et al.* 2015. Briefly, leaflet outlines were thresholded using ImageJ (Abramoff *et al.* 2004) and converted to.bmp files for analysis in SHAPE (Iwata & Ukai 2002), where each leaflet was converted into chaincode, oriented, and decomposed into harmonic coefficients. The harmonic coefficients were then converted into a data frame format and read into R (R Core Team 2018). The Momocs package (Bonhomme *et al.* 2014) was used to visualize mean leaflet shapes from each genotype/light treatment combination. The prcomp() function was used to perform a Principal Component Analysis (PCA) on only A and D harmonics so that only symmetric (rather than asymmetric) shape variance was considered (Iwata *et al.* 1998). The results were visualized using ggplot2 (Wickham 2016).

### Light microscopy analyses

The fully expanded fifth leaf was cleared with 95% methanol for 48h followed by 100% lactic acid. Stomatal pore area index (SPI) was calculated as (guard cell length)^2^ × stomatal density for the adaxial and abaxial epidermes and then added up (Sack *et al.* 2003). Stomatal density was calculated as number of stomata per unit leaf area, stomatal index as the proportion of guard cells to total epidermal cells. Minor vein density was measured as length of minor veins (<0.05 μm diameter) per unit leaf area.

For cross-sectional analyses, samples were collected from the medial region of the fully expanded fifth leaf and fixed in 70% formalin-acetic acid-alcohol (FAA) solution for 48h and then stored in 70% (v/v) aqueous ethanol. The samples were embedded in historesin (Leica Microsystems, Wetzlar, Germany), cut into cross-sections (5μm) with an automated rotary microtome (RM2155, Leica Microsystems, Wetzlar, Germany) and sequentially stained with toluidine blue. Images obtained in a light microscope (Zeiss, Axioscope A1 model, Thornwood, NY, USA) with attached Axiovision^®^ 105 color image capture system. Anatomical parameters were quantified using Image Pro-Plus^®^ software (version 4.5, Media Cybernetics, Silver Spring, USA).

Mesophyll surface area exposed to intercellular air spaces per leaf area (*S*_*mes*_*/S*) was calculated separately for spongy and palisade tissues as described by Evans et al. (1994). To convert the length in cross-sections to the surface area, a curvature correction factor was measured and calculated for each treatment according to Thain (1983) for palisade and spongy cells by measuring their width and height and calculating an average width/height ratio. The curvature factor correction ranged from 1.17 to 1.27 for spongy cells and from 1.38 to 1.45 for palisade cells. All parameters were analysed at least in four different fields of view. *S*_*m*_*/S* was calculated as an weighted average based on tissue volume fractions.

### Anatomical estimation of mesophyll conductance (g_m_)

The one-dimensional gas diffusion model of Niinemets & Reichstein (2003) as applied by Tosens et al. (2012) was employed to estimate the share of different leaf anatomical characteristics in determining mesophyll conductance (*g*_m_). *g*_m_ as a composite conductance for within-leaf gas and liquid components is given by:

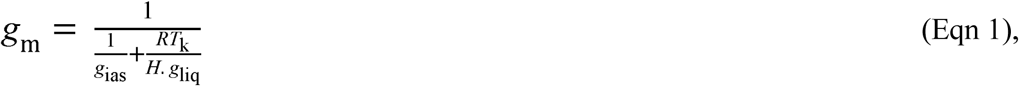

where *g*_ias_ is the gas phase conductance inside the leaf from substomatal cavities to outer surface of cell walls, *g*_liq_ is the conductance in liquid and lipid phases from outer surface of cell walls to chloroplasts, *R* is the gas constant (8.314 Pa m^3^ K^−1^ mol^−1^), *T*_k_ is the absolute temperature (K), and *H* is the Henry’s law constant (2938.4 Pa m^3^ mol^−1^). *g*_m_ is defined as a gas-phase conductance, and thus *H*/(*RT*_k_), the dimensionless form of Henry’s law constant, is needed to convert *g*_liq_ to corresponding gas-phase equivalent conductance (Niinemets & Reichstein 2003). In the model, the gas-phase conductance (and the reciprocal term, *r*_ias_) is determined by average gas-phase thickness, Δ*L*_ias_, and gas-phase porosity, *ƒ*_ias_ (fraction of leaf air space):

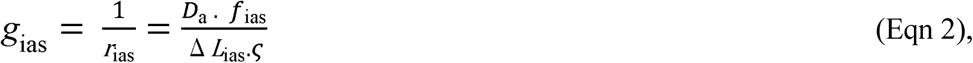

where *ς* is the diffusion path tortuosity (1.57 m m^−1^, value taken from Niinemets & Reichstein (2003) and *D*_a_ (m^2^ s^−1^) is the diffusion coefficient for CO_2_ in the gas phase (1.51 x 10^−5^ at 25 °C). Δ*L*_ias_ was taken as half the mesophyll thickness.

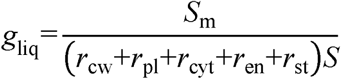

The term *r*_i_, where *i* stands either for cell wall (cw), plasma membrane (pl), cytosol (cyt), chloroplast envelope (en), and stroma (st) resistances are the partial determinants of the liquid-phase diffusion pathway. Cell wall thickness is the main determinant of liquid-phase resistance, and, as we found little variation for this parameter when comparing two studies conducted under different conditions (Berghuijs *et al.* 2015; Eid Gamel, Elsayed, Bashasha & Haroun 2016) we used the partial determinants of the liquid-phase diffusion pathway described in Berghuijs et al. (2015). In addition, *S*_*mes*_*/S*, a major determinant of *g*_liq_, was measured in this study. Total liquid-phase diffusion was scaled by the *S*_mes_/*S* as there was little cell wall area free of chloroplasts (Figure S3) reflecting a ratio between chloroplast and mesophyll area exposed to intercellular airspaces (*S*_c_/*S*_mes_) very close to 1.0 as also observed by Galmés et al. (2013).

### Carbon isotope composition

The fully expanded fifth leaf of five plants per treatment were harvested and ground to fine powder. Samples were sent to the Laboratory of Stable Isotopes (CENA, USP, Piracicaba, Brazil), where they were analysed for ^13^C/^12^C ratio using a mass spectrometer coupled to a Dumas elemental analyser ANCA-SL (Europa Scientific, Crewe, UK). Carbon isotope ratios were obtained in *δ*-notation, where

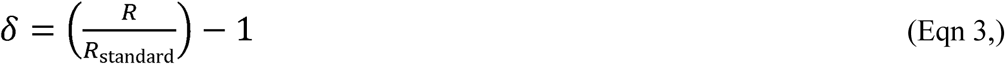

and *R* and *R*_standard_ are the isotope ratios of the plant sample and the Vienna Pee Dee Belemnite (VPDB) standard, respectively. *δ*^13^C of atmospheric CO_2_ was assumed to be −8 per mil. The *δ*^13^C values for the samples were then converted to carbon isotopic discrimination values, Δ^13^C = (*δ*_a_- *δ*_p_)/(1 + *δ*_p_), where *δ*_a_ is the *δ*^13^C of atmospheric CO_2_ and *δ*_p_ the *δ*^13^C of the plant material (Farquhar and Sharkey 1982).

### Gas exchange and chlorophyll fluorescence determinations

Gas exchange analyses were performed in MT and M82 plants at 40 and 50 days after germination, respectively. Gas exchange measurements were performed using an open-flow gas exchange system infrared gas analyzer (IRGA) model LI-6400XT (LI-Cor, Lincoln, NE, USA). The analyses were performed under common conditions for photon flux density (1000 μmol m^−2^s^−1^, from standard LiCor LED source), leaf temperature (25 ± 0.5°C), leaf-to-air vapor pressure difference (16.0 ± 3.0 mbar), air flow rate into the chamber (500 μmol s^−1^) and reference CO_2_ concentration of 400 ppm (injected from a cartridge), using an area of 2 cm^2^ in the leaf chamber. For dark respiration (R_d_) determination, plants were adapted to the dark at least 1h before the measurements, as described by Niinemets *et al* 2006.

Photochemical efficiency of photosystem II (φPSII) was determined by measuring the steady-state fluorescence (*F*_s_) and the maximum fluorescence (*F*_m_’), using a pulse of saturating light of approximately 8000 μmol photons m^−2^ s^−1^, as described by Genty *et al.* 1989. Photosynthetic light response curves were measured under ambient O_2_, with reference CO_2_ set to 400 μmol mol^−1^. After allowing full photosynthetic induction for 30-45 min, *A* was determined at *PPFD* steps 1500, 1200, 1000, 800, 600, 400, 300, 200, 150, 75, 50, 0 μmol m^−2^ s^−1^ at ambient temperature (25°C) and CO_2_ concentration (400 μmol mol^−1^) The light saturation point (*I*_s_), light compensation point (*I*_c_), light saturation CO_2_ assimilation rate (*A*_PPFD_) and the light utilization (1/Φ). *A/C*_i_ curves were constructed with step changes (50, 100, 150, 250, 400, 500, 700, 900, 1200, 1300, 1400 and 1600 μmol mol^−1^) of [CO_2_] under 1000 μmol m^−2^ s^−1^ light, at 25°C under ambient O_2_ supply. The maximum rate of carboxylation (*V*_*c*max_), maximum rate of carboxylation limited by electron transport (*J*_max_) and triose-phosphate utilization (TPU) were estimated by fitting the mechanistic model of CO_2_ assimilation proposed by Farquhar *et al.*1980. Additionally, *g*_m_ was tentatively estimated using the Ethier and Livingston (2004) method, which is based on fitting *A/C*_i_ curves with a non-rectangular hyperbola version of the FvCB which incorporates *g*_m_ in the model. Corrections for the leakage of CO_2_ into and out of the leaf chamber of the LI-6400 were applied to all gas-exchange data as described by Rodeghiero *et al.* 2007 using a *K*_CO2_ estimated as 0.4 μmol s^−1^

### Water relations

Leaf (Ψ_L_) or xylem (Ψ_X_) water potential were measured in the central leaflet of the fifth fully expanded leaf in MT and M82 plants 40 and 50 days of age, respectively, using a Scholander-type pressure chamber (model 1000, PMS Instruments, Albany, NY, USA). Ψ_L_ was measured in transpiring leaves, whereas Ψ_X_ was obtained from non-transpiring leaflets, assumed to be in equilibrium with the petiole water potential. The non-transpiring leaflet consisted of the lateral leaflet of the same leaf, which was covered with plastic film and foil the night before the measurements. Apparent hydraulic conductance (*K*_leaf_) were estimated using the transpiration rates and the water potential difference between the transpiring and non-transpiring leaflet according to Ohm’s law:

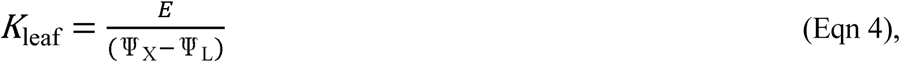

Where: *E* is the transpiration rate (mmol m^−2^ s^−1^) determined during gas exchange measurements, and (Ψ_L_ - Ψ_X_) corresponds to the pressure gradient between the transpiring and non-transpiring leaflet (MPa). Water potential and hydraulic conductance measurements were performed immediately after gas exchange analysis.

### Biochemical determinations

Biochemical analyses of the leaves were performed in MT and M82 plants 40 and 50 days after germination, respectively. The terminal leaflet of the sixth fully expanded leaf was collected around midday on a cloudless day, instantly frozen in liquid N_2_ and stored at −80°C. Subsequently, the samples were lyophilized at −48°C and macerated with the aid of metal beads in a Mini-Beadbeater-96 type cell disrupter (Biospec Products, Bartlesville, OK, USA). A 10-mg sample of ground tissue was added to pure methanol (700 μL), and the mixture was incubated at 70 °C for 30 min followed by a centrifugation (16,200 × *g*, 5 min). Supernatant was placed in new tubes in which was added chloroform and ultrapure water (375 μL and 600 μL, respectively). After new centrifugation (10,000 × *g*, 10 min), the concentrations of hexoses (glucose plus fructose) and sucrose were determined in the aqueous phase by a three step reaction in which hexokinase, phosphoglucose isomerase and invertase (Sigma Aldrich) were subsequently added to a reaction buffer containing ATP, NADH and glucose dehydrogenase (Sigma Aldrich) according to Fernie et al. (2001). The methanol-insoluble pellet was resuspended by adding 1mL 0.2M KOH followed by incubation at 95 °C. The resulted solution was used for subsequent protein quantification (Bradford method). Finally, 2M acetic acid was added (160 μL) to the resuspended pellet from which was quantified Starch by adding hexokinase in a buffer reaction as previously describe for sugars. Noteworthy, the above described protocol was previously detailed by Praxedes, DaMatta, Loureiro, G. Ferrão & Cordeiro 2006 and Ronchi *et al.* 2006 and includes some of the recommendations described by Quentin *et al.* 2015, such as the use of amyloglucosidase for starch extraction and the use of glucose and starch standards. Photosynthetic pigments (chlorophyll (a + b) content and carotenoids) were determined in the methanolic extract using the equations described in Porra et al. (1989) using a microplate reader.

### Agronomic parameters (yield and Brix)

The number of fruits per plant was obtained from fruit counts and the frequency of green and mature fruits was also determined separately. Fruit average weight was determined after individual weighing of each fruit, using a semi analytical balance with a sensitivity of 0.01 g (AUY220, Shimadzu, Kyoto, Japan). Yield per plant corresponds to the total weight of fruits per plant. The determination of the soluble solids content (°Brix, which is the % of soluble solids by weight) in the fruits was measured with a digital temperature-compensated refractometer, model RTD 45 (Instrutherm^®^, São Paulo, Brazil). Six ripe fruits per plant were evaluated in five replicates per genotype.

### Statistical analysis

The data were subjected to analysis of variance (ANOVA) using Assistat version 7.6 (http://assistat.com) and the means were compared by the Tukey test at the 5% level of significance (*P*≤0.05).

## Results

This study was performed on two tomato genetic backgrounds, cultivars Micro-Tom (MT) and M82, and their respective *obscuravenosa* (*obv*) mutant near-isogenic lines (NILs). First, we conducted a microscopic analysis of terminal leaflet cross-sections to confirm that, like all wild-type tomatoes and its wild relatives, MT harbors bundle sheath extensions (BSEs) in primary (*i.e.* midrib) and secondary veins of fully-expanded leaves (Fig. 1a). The *obv* mutant, on the other hand, lacks these structures, so that the veins appear obscure (hence the name of the genotype) (Fig. 1b). Chlorophyll fluorescence imaging revealed that this optical effect is due to the continuity of the palisade mesophyll on the adaxial side, and of the spongy mesophyll on the abaxial side in *obv*, which are both interrupted by BSEs in MT (Fig. 1c,d). The BSEs protrude toward the adaxial epidermis as columns of possibly parenchymatic or collenchymatic cells with thickened cell walls, whereas they thicken downward and are broadly based upon the lower epidermis (Fig. 1e-h). We next conducted a water + dye infiltration assay in the lamina, proving that, under similar pressure, intercellular spaces of the *obv* mutant were flooded almost twice (86.1% vs 47.3%, *P*=0.012) as much as for MT (Fig. S1). Dry patches were observed in MT, which shows that the presence of BSEs in secondary veins creates physically isolated compartments in the lamina (Fig. S1). We therefore follow the established nomenclature of ‘heterobaric’ for MT and ‘homobaric’ for *obv*.

**Fig. 1.**
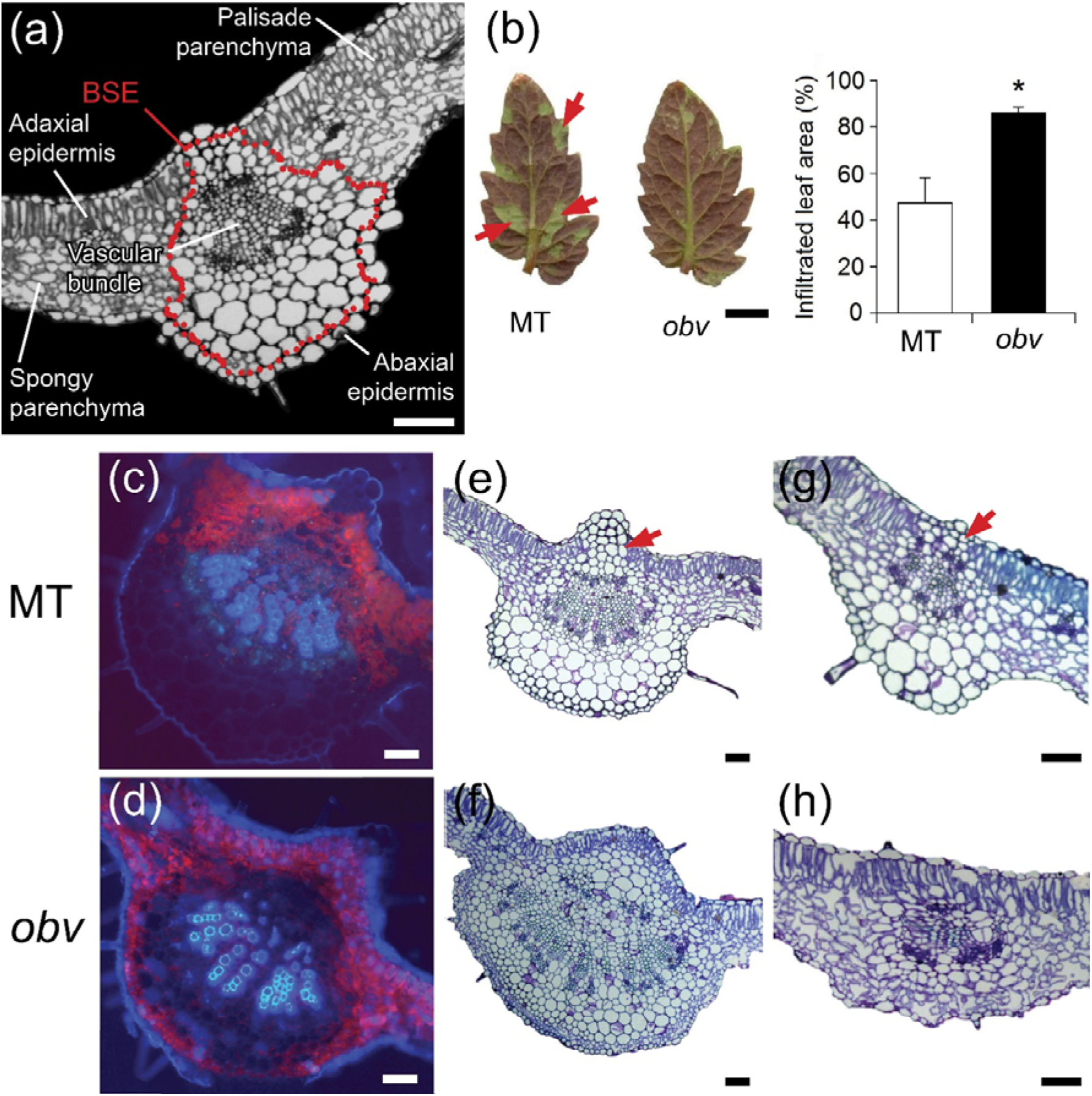
Leaf anatomical differences between Micro-Tom (MT) and the *obscuravenosa* (*obv*) mutant. (a) Semi-schematic representation of cross-sectional anatomy of a wild-type (MT) secondary vein. BSE= bundle sheath extension. (b) Representative images of terminal leaflets from fully expanded leaves infiltrated with 1% fuchsin acid solution applying 0.027 MPa of pressure during 2 minutes showing dry patches (arrowheads) in MT, as opposed to uniform infiltration in *obv*. Scale bar=1 cm. Bars are mean values ± s.e.m. (n=4). Asterisk indicates significant difference by Student’s t-test (*P* <0.05) (c) Chlorophyll fluorescence showing interruption of the palisade mesophyll on the adaxial side, and of the spongy mesophyll on the abaxial side by BSE cells in MT, which are absent in *obv* (d). (e-h) Cross-sections of the leaf lamina at the midrib (e,f) and a secondary vein (g,h) show the presence (MT) and absence (*obv*) of bundle sheath extensions (BSEs). The BSEs have a columnar nature protruding toward the adaxial epidermis (arrowheads), with thickened cells walls, whereas they thicken downward and are broadly based upon the lower epidermis. Scale bars= 1 cm (leaflets) and 100 μm (midrib and secondary vein).

### Irradiance level alters leaf shape and structural parameters differentially in heterobaric and homobaric leaves

We began by conducting an analysis of leaflet shape between the treatments. A Principal Component Analysis (PCA) on harmonic coefficients contributing to symmetric shape variation separates MT and *obv* genotypes, but failed to show large differences in shape attributable to light treatment (Fig 2a). To visualize the effects of genotype and light, we superimposed mean leaflet shapes from each genotype-light combination (Fig 2b). *obv* imparts a wider leaflet shape relative to MT, regardless of light treatment. Light treatment did not discernibly affect leaflet shape.

**Fig. 2.**
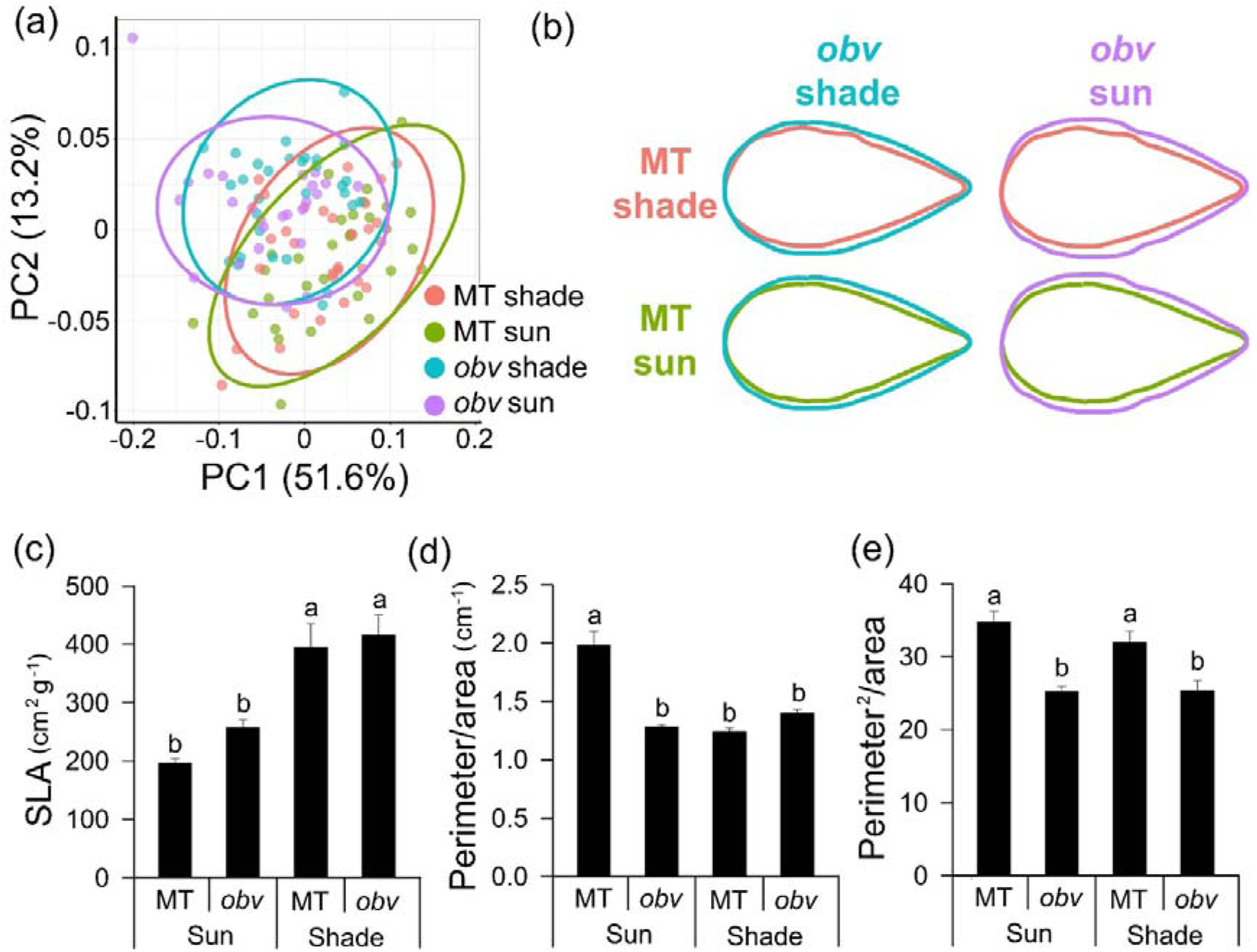
Irradiance level differentially alters morphology in heterobaric and homobaric leaves. (a) Principal Component Analysis (PCA) on A and D harmonic coefficients from an Elliptical Fourier Descriptor (EFD) analysis shows distinct symmetric shape differences between MT and *obv* leaflets, but small differences due to light treatment. 95% confidence ellipses are provided for each genotype and light treatment combination, indicated by color. (b) Mean leaflet shapes for MT and *obv* in each light treatment. Mean leaflet shapes are superimposed for comparison. Note the wider *obv* leaflet compared to MT. MT shade, red; MT sun, green; *obv* shade, blue; *obv* sun, purple. (c) Specific leaf area (SLA); (d-e) relationship between perimeter/area and perimeter^2^/area. Bars are mean values ± s.e.m. (n=5). Different letters indicate significant differences by Tukey’s test at 5% probability.

Sun leaves had reduced total specific leaf area (SLA) compared to shade leaves in both MT and the *obv* mutant (Fig 2c). Shading increased SLA values by 101% and 62% for MT and *obv* plants, respectively, when compared to plants in the sun treatment. Terminal leaflets of fully expanded MT sun leaves had 62% higher perimeter/area than MT shade leaves, unlike *obv* where we found no difference between irradiance levels (Fig 2d). Perimeter^2^/area, which, unlike perimeter/area is a dimensionless measure of leaf shape (and, therefore, does not inherently scale with size), was strongly dependent on genotype and not influenced by irradiance (Fig 2e).

### Growth irradiance alters leaf hydraulic conductance in heterobaric but not in homobaric leaves in different tomato genetic backgrounds

Leaf hydraulic conductance (*K*_leaf_) is a key parameter determining plant water relations, as it usually scales up to the whole plant level (Sack & Holbrook 2006). Shading decreased *K*_leaf_ in the heterobaric genotype: MT shade leaves had 41% lower *K*_leaf_ than sun leaves (14.95 ± 1.91 *vs* 25.36 ± 1.32 mmol H_2_O m^−2^ s^−1^ MPa^−1^) (Fig 3a,b). Homobaric and heterobaric leaves in the M82 tomato background (Fig 3c), showed a similar leaf vein phenotype as in MT (Fig 3d) showed consistently similar results, where shade leaves had 36% lower *K*_leaf_ than sun leaves (18.72 ± 0.59 *vs* 29.6 ± 2.1 mmol H_2_O m^−2^ s^−1^ MPa^−1^) (Fig 3d). The *obv* mutant, on the other hand, showed similarly low *K*_leaf_ values in either condition and in both genetic backgrounds (MT sun: 17.86 ± 1.26 *vs* shade: 17.87 ± 2.14 mmol H_2_O m^−2^ s^−1^ MPa^−1^; M82: sun: 19.19 ± 2.24 *vs* shade: 19.17 ± 2.67 mmol H_2_O m^−2^ s^−1^MPa^−1^) (Fig 3b,e). The results were consistent between tomato backgrounds, even though both cultivars differ markedly in leaf lamina size and other leaf structural parameters.

**Fig. 3.**
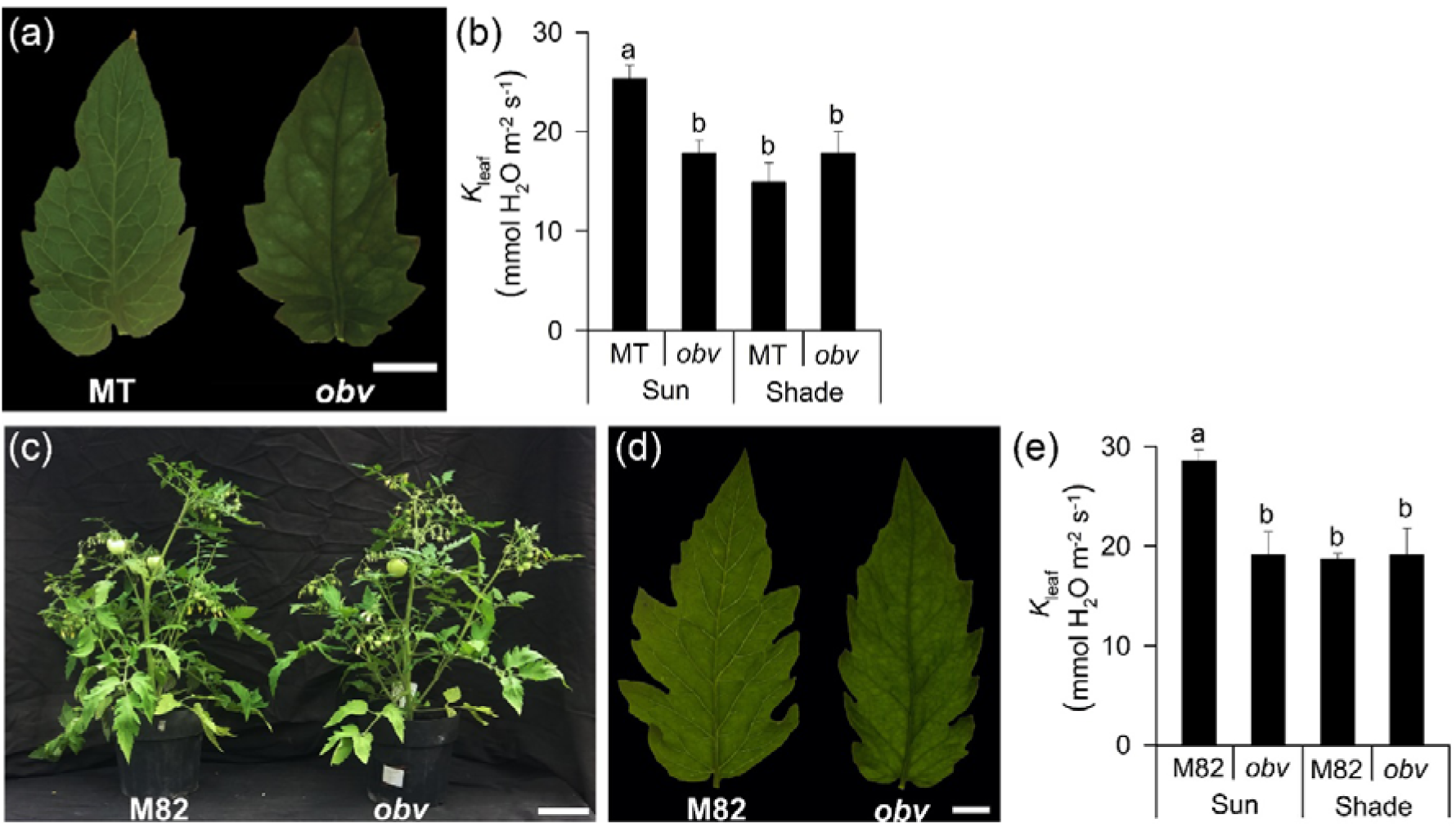
(a) Representative terminal leaflets of tomato cv Micro-Tom (MT, heterobaric) and the *obscuravenosa* mutant (*obv*, homobaric) leaves, showing translucent and dark veins, respectively. Bar=1cm. (b) Leaf hydraulic conductance (*K*_leaf_) in homobaric and heterobaric leaves grown in either sun or shade conditions. Bars are mean values ± s.e.m. (n=3). Different letters indicate significant differences by Tukey’s test at 5% probability. (c) Representative F1 plants and (b) terminal leaflets of Micro-Tom × M82 (M82, heterobaric) and Micro-Tom *obv* × M82 (*obv*, homobaric). Scale bars= 10 cm (c) and 1 cm (d). (e) *K*_leaf_ in F_1_ plants of M82 × MT (M82, heterobaric) and F_1_ plants of M82 × MT-*obv*, (*obv*, homobaric) leaves from plants grown in either sun or shade conditions. Bars are mean values ± s.e.m. (n=5). Different letters indicate significant differences by Tukey’s (*P* <0.05).

### Shading reduces stomatal conductance in heterobaric leaves, whereas homobaric leaves maintain similarly low values under both conditions

Previous work has suggested that BSEs could influence photosynthetic assimilation rate (*A*) by increasing light transmission within the mesophyll (Karabourniotis *et al.* 2000). To ascertain whether this was the case in our genotypes, we determined photosynthetic light response curves on fully expanded terminal leaflets attached to plants growing in the greenhouse under sun or shade treatments (Fig S1). Although no statistical differences were found in the light response of *A* between heterobaric MT and homobaric *obv* plants (Fig S1), the light saturation point (*I*_s_) was lower in shade *obv* than in the other treatments (Table S1)

Since the presence of BSEs can affect lateral flow of CO_2_ within the leaf blade (Pieruschka *et al.* 2006; Morison *et al.* 2007), we next analysed the response of *A* to varying internal partial pressure of CO_2_ in the substomatal cavity (*C*_i_) (Table 2). The apparent maximum carboxylation rate of Rubisco (*V*_cmax_), the maximum potential rate of electron transport in the regeneration of RuBP (*J*_max_) and the speed of use of triose-phosphates (TPU) were reduced by 20.0%, 20.2% and 21.1%, respectively, for shade compared to sun MT plants. In *obv*, the respective drop between sun and shade plants the same parameters was 10.0%, 7.0% and 6.0% respectively (Table 2).

**Table 2.** Gas exchange parameters determined in fully-expanded leaves of heterobaric (Micro-Tom, MT) and homobaric (*obscuravenosa*, *obv*) in two irradiance levels (sun/shade, 900/300 μmol photons m^−2^ s^−1^).

|  | Sun |  | Shade |  |
| --- | --- | --- | --- | --- |
|  | MT | *obv* | MT | *obv* |
| $A$ ($\mu\text{mol CO}_2 \text{ m}^{-2} \text{s}^{-1}$) | $21.29 \pm 1.34\text{a}$ | $20.74 \pm 1.44\text{a}$ | $17.07 \pm 0.83\text{b}$ | $20.26 \pm 0.48\text{a}$ |
| $g_s$ ($\text{mol m}^{-2} \text{s}^{-1}$) | $0.373 \pm 0.039\text{a}$ | $0.275 \pm 0.020\text{b}$ | $0.263 \pm 0.016\text{b}$ | $0.278 \pm 0.018\text{b}$ |
| $\text{TE}_i$ ($A/g_s$) | $59.16 \pm 3.25\text{b}$ | $76.26 \pm 2.16\text{a}$ | $65.51 \pm 2.08\text{b}$ | $74.11 \pm 3.55\text{a}$ |
| $V_{c,\text{max}}$ ($\mu\text{mol m}^{-2} \text{s}^{-1}$) | $82.7 \pm 6.04\text{a}$ | $80.5 \pm 6.26\text{a}$ | $66.8 \pm 4.38\text{a}$ | $72.7 \pm 7.72\text{a}$ |
| $J_{\text{max}}$ ($\mu\text{mol m}^{-2} \text{s}^{-1}$) | $167.5 \pm 5.74\text{a}$ | $155.5 \pm 8.48\text{a}$ | $133.5 \pm 4.54\text{b}$ | $130.2 \pm 3.31\text{b}$ |
| TPU ($\mu\text{mol m}^{-2} \text{s}^{-1}$) | $12.1 \pm 0.34\text{a}$ | $11.0 \pm 0.62\text{a}$ | $9.6 \pm 0.36\text{b}$ | $10.3 \pm 0.1\text{a}$ |
| $R_d$ ($\mu\text{mol CO}_2 \text{ m}^{-2} \text{s}^{-1}$) | $1.49 \pm 0.43 \text{ a}$ | $1.80 \pm 0.45 \text{ a}$ | $1.42 \pm 0.38 \text{ a}$ | $1.45 \pm 0.39 \text{ a}$ |

Values are means ± s.e.m (n=8 for $A$ , $g_s$ and $\text{TE}_i$ ; n=6 for other parameters). Values followed by the same letter in each row were not significantly different by Tukey test at 5% probability.

The hyperbolic relationship between *A* and *g*_s_ measured at ambient CO_2_ was not altered by irradiance level (Fig 4a, b). The lower limit for *g*_s_ values was remarkably similar between genotypes in both light conditions (~0.2 mol m^−2^ s^−1^). A 30% decrease in *g*_s_ with a concomitant limitation to *A* was observed in shade MT (Table 2),. In the *obv* mutant, *g*_s_ was lower in the sun (similar value to shade MT) and remained essentially unchanged by shading, as did *A*. The *A*/*g*_s_ ratio, or intrinsic water-use efficiency (WUE_i_), was therefore higher in homobaric *obv* plants than in heterobaric MT under both irradiance levels (Table 2). A similar, although not statistically significant difference (possibly owing to the lower number or replicates, n=5) was found in M82 (Fig S2). Dark respiration was not affected by genotype or irradiance level (Table 2). The chlorophyll fluorescence analyses revealed a higher quantum yield of photosystem II photochemistry (ΦPSII) and electron transport rate (ETR) in homobaric *obv* plants grown in the sun than in all other treatments. No differences between treatments were found in the photochemical and non-photochemical quenching (Table S4).

**Fig. 4.**
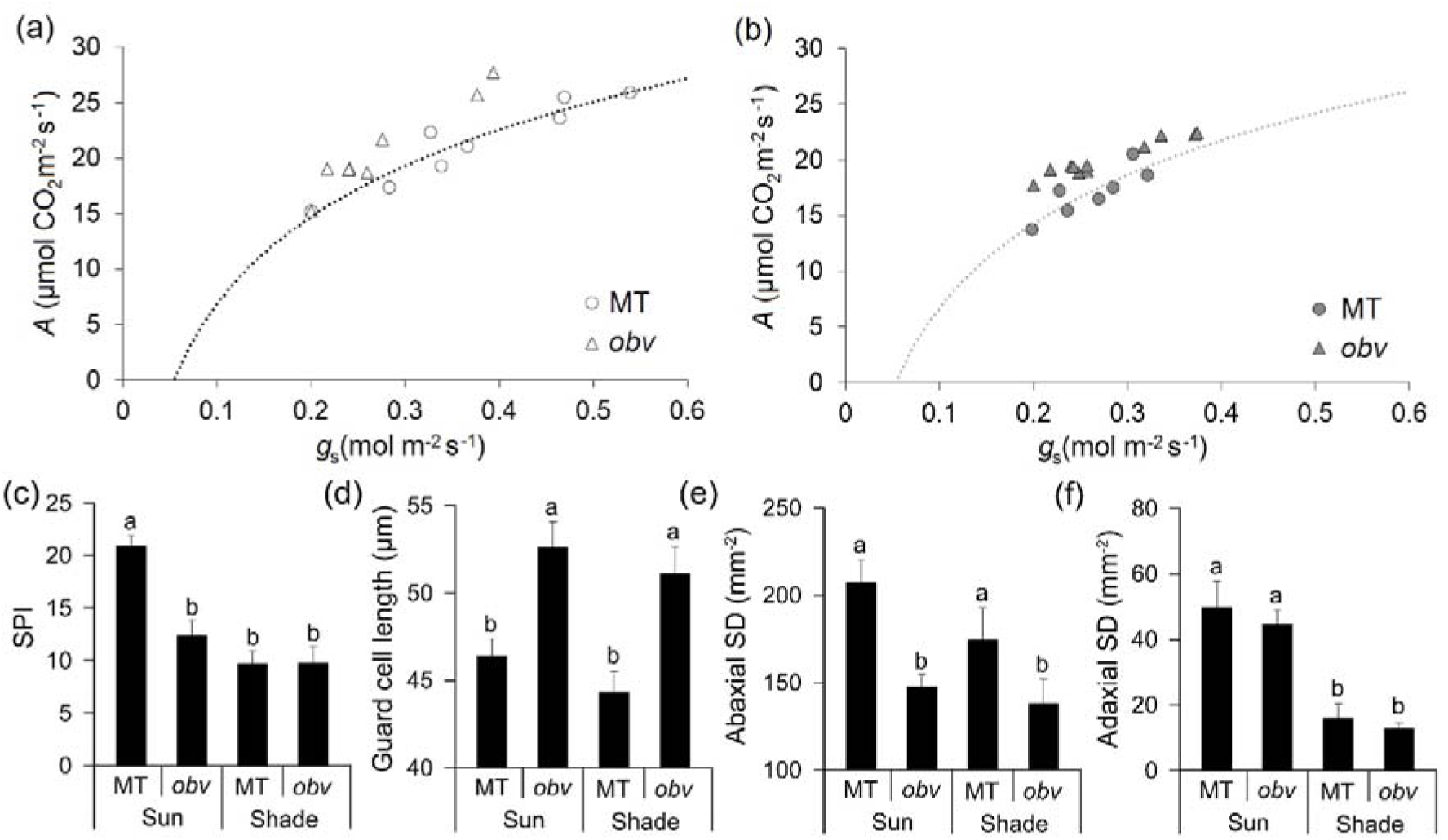
Homobaric leaves maintain lower stomatal conductance in both sun and shade conditions. Relationship between photosynthetic CO_2_ assimilation rate (*A*) and stomatal conductance (*g*_s_) for Micro-Tom (MT) and the *obscuravenosa* (*obv*) mutant plants grown in the sun (a) or shade (b). A rectangular hyperbolic function was fitted in each panel. Each point corresponds to an individual measurement carried out at common conditions in the leaf chamber: photon flux density (1000 μmol m^−2^ s^−1^, from an LED source), leaf temperature (25 ± 0.5°C), leaf-to-air vapor pressure difference (16.0 ± 3.0 mbar), air flow rate into the chamber (500 μmol s^−1^) and reference CO_2_ concentration of 400 ppm (injected from a cartridge). (c-f) Stomatal traits are differentially affected by irradiance in heterobaric and homobaric tomato leaves. (a) SPI: stomatal pore area index, calculated as (guard cell length)^2^ × stomatal density for the adaxial and abaxial epidermes and then added up; (b) Guard cell length; (c-d) Stomatal density (number of stomata per unit leaf area); Data shown as means ± s.e.m. (n=6). Different letters indicate significant differences by Tukey’s test at 5% probability.

### Stomatal Pore Area Index is altered by irradiance in heterobaric but not homobaric leaves

Stomatal conductance (*g*_s_) is influenced by the maximum stomatal conductance (*g*_max_), which is in turn determined by stomatal size and number (Parlange & Waggoner 1970; Franks & Beerling 2009). To further explore the basis for the differential *g*_s_ response to irradiance between genotypes, we analysed stomatal traits in terminal leaflets of fully expanded leaves. Stomatal pore area index (SPI, a combined dimensionless measure of the stomatal density and size) was increased only in MT sun leaves (Fig 4c), compared to all the other treatments‥ Guard cell length, which is linearly related to the assumed maximum stomatal pore radius, was greater in *obv* than in MT and was not affected by the irradiance levels (Fig 4d). Thus, the main driver of the difference in SPI was stomatal density, particularly on the abaxial side, which represents a quantitatively large contribution (Fig 4e). Adaxial stomatal density was reduced in the shade in both genotypes, with no differences between them within irradiance levels (Fig 4f).

### Minor vein density, modelled mesophyll conductance to CO_2_ and carbon isotope discrimination are differentially altered by irradiance levels in heterobaric and homobaric leaves

Leaf lamina thickness was reduced by shading in both genotypes, with no difference between them (Fig 5). These results are in good agreement with the reduced specific leaf area (SLA) in shade-grown plants (Fig 1c). The palisade to spongy mesophyll thickness ratio was increased by shading, independently of genotype (Fig 5c). Thickness of the abaxial epidermis, a proxy for stomatal depth, did not vary in MT between irradiance levels, but was reduced in shaded *obv* plants (Fig 5d). Intercellular air spaces in the lamina comprised close to 10% of the cross-sectional area in MT and *obv* plants grown in the sun, but when plants were grown in the shade, it was increased to 12% in MT and 17% in *obv* (Fig 5e). As venation is a key trait that influences water distribution in the lamina, we assessed minor vein density (tertiary and higher orders) and observed a genotype×irradiance interaction (Fig 5b). Vein density was reduced in both genotypes by shading, but more strongly in MT than in *obv* (Fig 5f).

**Fig. 5.**
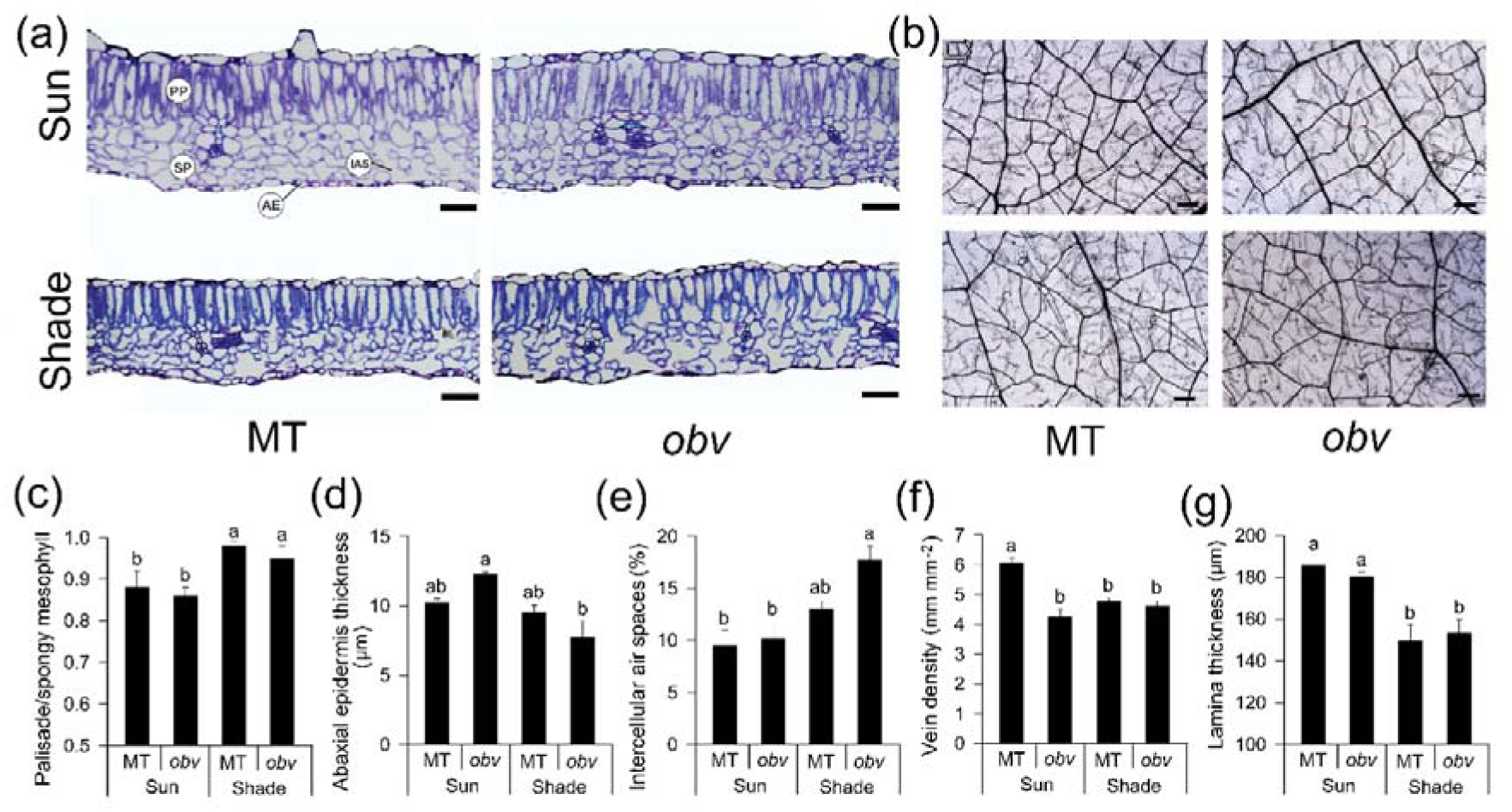
Irradiance level differentially alters leaf anatomical parameters in heterobaric and homobaric leaves. (a) Representative cross-sections of tomato cv Micro-Tom (MT, heterobaric) and the *obscuravenosa* mutant (*obv*, homobaric) leaves from plants grown in either sun or shade. The background was removed for clarity. PP: palisade parenchyma; SP: spongy parenchyma; IAS: intercellular air spaces; AE: abaxial epidermis. Scale bars=50 μm (b) Representative plates showing the pattern and density of minor veins in 7.8 mm^2^ sections in mature, cleared leaves. Scale bar=200 μm. (c-g) Histograms with mean values ± s.e.m. (n=6) for the ratio between palisade and spongy parenchyma thickness; thickness of the abaxial epidermis; the proportion of intercellular air spaces and the density of minor (quaternary and higher order) veins measured in cleared sections of the leaves and lamina thickness. Different letters indicate significant differences by Tukey’s test at 5% probability.

We next used anatomical data (Fig S3) to estimate mesophyll conductance to CO_2_ (*g*_m_), a key parameter linking leaf hydraulics, photosynthetic function and leaf anatomy (Flexas *et al.* 2013; Tomás *et al.* 2013). Our estimates suggest that the lack of BSEs significantly altered the value of *g*_m_ in response to shading, whereas the genotypes did not vary significantly for this parameter when grown in the sun (Table 3). As a way to validate our results, and also due to its intrinsic interest as a proxy for *C*_i_*/C*_a_ (the ratio of CO_2_ concentration inside and outside the leaf) (Condon *et al.* 2004), we next determined carbon isotope composition (*δ*^13^C) in leaves from the same plants used for the anatomy and gas exchange measurements (Table S2). The *obv* mutation had a differential effect on carbon isotope discrimination (Δ^13^C), a parameter that is linearly and negatively correlated to long-term water-use efficiency (WUE) of plants. Whereas the presence of the *obv* mutation increased Δ^13^C in the sun (thus, decreased WUE), it had the opposite effect in the shade [lower Δ^13^C and higher WUE (Fig S4)].

**Table 3.** Mesophyll conductance modeled from anatomical characteristics (*g*_m_anatomical_), gas phase conductance inside the leaf from substomatal cavities to outer surface of cell walls (*g*_ias_), conductance in liquid and lipid phases from outer surface of cell walls to chloroplasts (*g*_ias_) and mesophyll surface area exposed to intercellular airspace (*S*_m_/*S*) determined in fully-expanded leaves of heterobaric (Micro-Tom, MT) and homobaric (*obscuravenosa*, *obv*) in two irradiance levels (sun/shade, 900/300 μmol photons m^−2^ s^−1^).

|  | Sun |  | Shade |  |
| --- | --- | --- | --- | --- |
|  | MT | *obv* | MT | *obv* |
| $g_{m\_anatomical}$ ($\text{mol m}^{-2} \text{s}^{-1}$) | $0.107 \pm 0.005\text{c}$ | $0.132 \pm 0.005\text{b}$ | $0.124 \pm 0.006\text{bc}$ | $0.162 \pm 0.004\text{a}$ |
| $g_{ias}$ ($\text{mol m}^{-2} \text{s}^{-1}$) | $0.466 \pm 0.028\text{b}$ | $0.419 \pm 0.060\text{b}$ | $0.780 \pm 0.057\text{a}$ | $1.029 \pm 0.089\text{a}$ |
| $g_{liq}$ ($\text{mol m}^{-2} \text{s}^{-1}$) | $0.117 \pm 0.006\text{b}$ | $0.170 \pm 0.013\text{a}$ | $0.125 \pm 0.005\text{b}$ | $0.163 \pm 0.007\text{a}$ |
| $S_{mes}/S$ ($\text{m}^2 \text{m}^{-2}$) | $6.3 \pm 0.30\text{b}$ | $9.2 \pm 0.72\text{a}$ | $6.8 \pm 0.29\text{b}$ | $8.8 \pm 0.36\text{a}$ |

Values are means ± s.e.m (n=4). Values followed by the same letter in each row were not significantly different by Tukey test at 5% probability.

### Carbohydrate and pigment contents in heterobaric and homobaric leaves under different irradiance

We assessed a basic set of compounds related to primary cell metabolism in MT and *obv* under both sun and shade conditions, along with photosynthetic pigments (Table S3). As expected, carbohydrate concentrations were strongly influenced by irradiance level (Table S3). Shading promoted a decrease in starch content in both genotypes, but of a considerable greater magnitude in MT (−45.0%) than in *obv* (−28.5%) compared to sun plants (Table S3). Glucose and fructose were increased in the shade, with no difference between genotypes. The chlorophyll a/b ratio was similar for all plants. A slight increase in carotenoid levels was found in *obv* shade plants (Table S4).

### Morphological and physiological differences between heterobaric and homobaric plant grown under different irradiances do not affect dry mass accumulation or fruit yield

To determine whether the anatomical and physiological differences described above scale up to the whole-plant level and affect carbon economy and agronomic parameters of tomato, we determined dry mass and fruit yield in sun-and shade-grown plants of MT and *obv*. There was no difference in plant height or in the number of leaves before the first inflorescence, for plants of either genotype in both light intensities (Table 4). There was a decrease in stem diameter in shade MT and *obv* plants, compared to sun plants. Leaf insertion angle relative to the stem, however, was steeper in the *obv* mutant under both irradiance conditions. Different light intensities did not change leaf dry weight, *obv* plants showed a 24.3% reduction in stem dry weight, 46.4% in root dry weight and 31% in total dry weight when compared to the sun treatment. The results were similar for MT, so no changes in dry mass allocation pattern were discernible between genotypes. Side branching is one of the most common morphological parameters affected by shading (Casal 2013). A decrease in side branching was found in both genotypes upon shade treatment, with no differences between them (Fig S5).

**Table 4.** Plant morphological parameters evaluated 40 days after germination (dag) in heterobaric (Micro-Tom, MT) and homobaric (*obscuravenosa*, *obv*) tomatoes grown in two irradiance levels (sun/shade, 900/300 μmol photons m^−2^ s^−1^).

|  | Sun |  | Shade |  |
| --- | --- | --- | --- | --- |
|  | MT | *obv* | MT | *obv* |
| Plant height (cm) | 9.90 ± 0.30a | 10.53 ± 0.28a | 10.15 ± 0.62a | 10.63 ± 0.18a |
| Leaves to 1<sup>st</sup> inflorescence | 6.75 ± 0.25a | 6.50 ± 0.18a | 6.62 ± 0.18a | 6.75 ± 0.25a |
| Leaf insertion angle (°) | 82.8 ± 2.32a | 73.1 ± 3.50b | 81.8 ± 4.30a | 65.5 ± 3.72b |
| Stem diameter (cm) | 0.40 ± 0.02a | 0.38 ± 0.03a | 0.28 ± 0.01b | 0.28 ± 0.01b |
| *Dry weight (g)* |  |  |  |  |
| Leaves | 1.30 ± 0.17a | 1.35 ± 0.06a | 1.07 ± 0.11a | 1.05 ± 0.08a |
| Stem | 2.17 ± 0.14ab | 2.49 ± 0.19ab | 1.54 ± 0.18b | 1.72 ± 0.07a |
| Roots | 0.80 ± 0.06a | 0.80 ± 0.04a | 0.50 ± 0.03b | 0.43 ± 0.04b |
| Total | 4.28 ± 0.34ab | 4.65 ± 0.28a | 3.12 ± 0.32b | 3.21 ± 0.14b |

Dry weight was determined through destructive analysis in plants 65 dag (n = 5). Values are means ± s.e.m (n=6).
Values followed by the same letter were not significantly different by Tukey test at 5% probability.

Vegetative dry mass accumulation was affected solely by irradiance level with no influence of the genotype, and therefore, independent of the presence or absence of BSEs. To ensure that potential differences arising from altered partitioning or allocation of carbon were not overlooked, we also assessed reproductive traits, *i.e.* parameters related to tomato fruit yield. Average fruit yield per plant was reduced by shading, but did not differ between genotypes within each irradiance condition, in two different tomato genetic backgrounds (MT and M82) (Table S5). The content of soluble solids in the fruit (°Brix), a parameter of agronomic interest, was also consistently stable across genotypes and treatments.

## Discussion

Heterobaric or homobaric plants are defined based on the presence or absence of BSEs, a structural characteristic associated with certain life forms and ecological distribution. Most of the studies addressing the function of BSEs have been based on large-scale multi-species comparisons, which restricts the conclusions to a statistical effect. Many structural, photosynthetic and hydraulic leaf traits are strongly co-ordinated and co-selected, therefore reducing the discriminating power of analyses involving species of different life forms and ecological background (Lloyd *et al.* 2013). Here, we compared different genotypes of a single herbaceous species (tomato) varying for a defined and ecologically relevant leaf structural feature: the presence of BSEs. The *obv* mutant lacks BSEs and thus produces homobaric leaves, compared to tomato cultivar Micro-Tom (MT), that has heterobaric leaves (Zsögön *et al.* 2015). We cultivated the plants under contrasting levels of irradiance (sun vs shade) and investigated leaf structure, hydraulics and photosynthetic function. We hypothesized that homobaric leaves, lacking a key physical feature that increases carbon assimilation and leaf hydraulic integration, would exhibit less plasticity than heterobaric leaves in their response to environmental conditions.

The presence or absence of BSEs did not affect general leaf morphology in either sun or shade conditions. SLA and leaf shape were altered by irradiance level, but without differences between genotypes. A generally higher photosynthetic capacity has been described for heterobaric species (Inoue *et al.* 2015), partially attributed to the optical properties of BSEs that enhance light transmission within the leaf mesophyll (Karabourniotis *et al.* 2000; Nikolopoulos *et al.* 2002). We did not observe such a photosynthetic advantage for heterobaric plants grown in high irradiance, but rather similar *A* values for both genotypes; indeed, the only difference we found for this genotype was a higher *g*_s_ which, despite not conferring higher *A*, might be beneficial in terms of latent heat loss, resulting in an improved thermal balance. Shading, on the other hand, reduced *A* in heterobaric MT plants, but not in *obv*. Since *g*_s_ and *V*_cmax_ were identical for both treatments, a higher *A* could be explained by a higher *g*_m_, and, consequently, higher chloroplast CO_2_ concentration.

We found that *obv* plants in the shade presented a high amount of intercellular air spaces and a high mesophyll surface area exposed to the intercellular air spaces (*S*_mes_/*S*). It seems that the absence of BSEs led to a higher *S*_mes_/*S* as they allowed more space to become available between palisade cells; on the other hand, presence of BSEs would ‘push’ palisade cells against each other, decreasing their exposure to the intercellular air spaces. An expected outcome of a higher *S*_mes_/*S* is to increase the anatomical *g*_m_, as it was the case for the *obv* plants in the shade (Table S6). However, our alternative *g*_m_ estimate (using the Ethier method, which takes into account both anatomical and biochemical *g*_m_ components) did not indicate any difference among plants (Table S6). Such discrepancy between the different estimates might reside in an important contribution from the biochemical components of *g*_m_ which is believed to be influenced by carbonic anhydrase and aquaporins expression (Flexas, Ribas-Carbó, Diaz-Espejo, Galmés & Medrano 2008; Tomás *et al.* 2013). In any case, our findings points to the need of further investigation of the role of BSEs on *g*_m_ using more refined methodologies (Pons *et al.* 2009).

In the shade, *g*_s_ was not changed between genotypes, thus resulting in an enhanced ratio between photosynthetic carbon gain and transpiratory water loss in homobaric *obv* plants. This observation was borne out by the reduced Δ^13^C in *obv* compared to the heterobaric MT. Long-term WUE is therefore higher in homobaric plants than in heterobaric plants in the shade, whereas the opposite is true in sun conditions. This provides a reasonable working hypothesis to explain the strongly biased ecological distribution of hetero-and homobaric species.

The higher incidence of heterobaric species in the canopy of both temperate and tropical forest canopies has been attributed to the effect of BSEs on leaf hydraulic integration (Kenzo *et al.* 2007; Inoue *et al.* 2015; Kawai *et al.* 2017). *K*_leaf_ was higher in heterobaric than in homobaric sun plants, consistent with the notion that BSEs act as an additional extra-xylematic pathway for the flow of liquid water thus enabling the maintenance of a higher *g*_s_ (Zwieniecki *et al.* 2007; Buckley *et al.* 2011). On the other hand, homobaric and heterobaric leaves showed similar *K*_leaf_ values in the shade, indicating that the presence of BSEs differentially affects leaf hydraulic architecture in response to irradiance. *K*_leaf_ is dynamically influenced by irradiance over different time scales, in the short-term by yet unknown factors (Scoffoni *et al.* 2008), and in the long-term by developmental plasticity altering leaf structural and physiological traits (Scoffoni *et al.* 2015). Buckley *et al.* (2015) found that BSEs increased *K*_leaf_ by 10%. They found that heterobaric species had 34% higher *K*_leaf_, but this must have been due to traits other than BSEs themselves. Interestingly, under high irradiance (sun), *K*_leaf_ was *c*. 30% higher in MT in comparison to *obv* plants, which is in line with the Buckley *et al.* (2015) estimates. A possible role for aquaporins present in the BS and/or the mesophyll has been proposed (Cochard *et al.* 2007) and it is known that aquaporins have their expression reduced under shade (Laur and Hacke 2013). Thus, it seems reasonable to assume that other *K*_leaf_ components were downregulated under shade, masking the contribution of BSEs to *K*_leaf_.

A large set of leaf physiological and structural traits shift in tandem in response to irradiance (Scoffoni *et al.* 2015). Particularly, plants developing under high light present a higher thermal energy load, which is dissipated mainly through leaf transpiration (Martins *et al.* 2014). In order to achieve higher transpiration rates, hydraulic supply and demand must be balanced, and vein density and patterning is coordinated with stomatal distribution to optimize resource utilization (Brodribb and Jordan 2011). Such coordination occurs across vascular plant species, but exactly how veins and stomata “communicate” with each other remains to be elucidated (Carins Murphy *et al.* 2017). In this sense, one of the proposed roles of BSEs is to act as a hydraulic linkage route between the vascular bundles and the epidermis, integrating these otherwise separated tissues (Zwieniecki *et al.* 2007). Here, we found that the presence of BSEs allowed a highly plastic coordination between veins and stomata, upregulating hydraulic supply and demand under high light (Fig 6). On the other hand, in genotypes lacking BSEs, the abaxial stomata and vein densities remained unchanged (Fig 6). At the moment there is no evidence to suggest BSEs are directly responsible for the plasticity in VLA and stomatal pore area index, nor that if they are responsible, it is because of an hydraulic effect on stomata. That seems unlikely, given that stomatal patterning mostly takes place before leaves begin to expand and transpire substantially. More data is needed to address this point. Another potential structural benefit of BSEs would be the provision of mechanical support, acting analogously to a suspension bridge, partially relieving the vein system from such duty and allowing heterobaric leaves greater flexibility in vein spacing compared to homobaric ones. Thus, the presence of BSEs could represent a hub coordinating trait plasticity in response to irradiance.

**Fig. 6.**
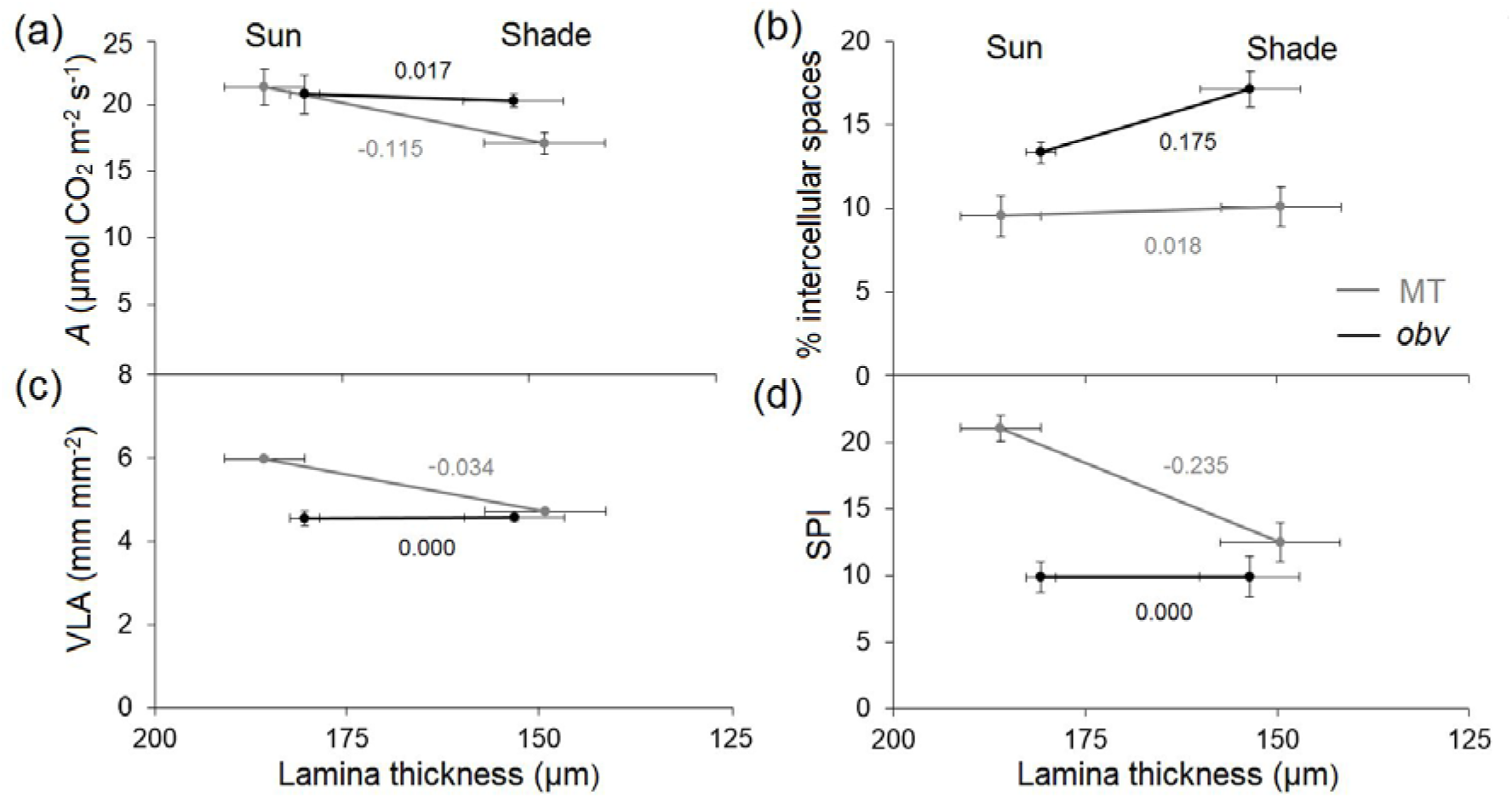
Reaction norms of structural and physiological traits in relation to leaf thickness in two irradiance levels in homobaric and heterobaric leaves. (a) light-saturated photosynthetic assimilation rate (*A*); (b) proportion of intercellular air spaces in the lamina, (c) minor vein per unit leaf area (VLA) and (d) stomatal pore area index (adimensional). The values of the slopes are shown next to each line. Error bars are s.e.m.

An open question is why the structural and physiological effects of the absence of BSEs in a leaf do not scale up to whole-plant carbon economy and growth. In other words, under what set of environmental conditions (if there is one) does the presence or absence of BSEs result in a significant fitness (*i.e.* survival and reproduction) difference between genotypes? The *obv* mutation has been incorporated by breeders in many tomato cultivar and hybrids (Jones *et al.* 2007), suggesting that it can confer some agronomic advantage. The present work was limited to analyzing the effect of discrete differences in irradiance and thus represents only a starting point to answering this question. The strong plasticity of plant development in response to irradiance (all other conditions being similar) could be the reason why potential economic differences between genotypes were canceled out within a given light environment. It is not possible to rule out that stronger quantitative differences in irradiance levels other than the ones tested here could tilt the phenotypic and fitness scales in favor of one of the leaf designs (*i.e.* heterobaric/homobaric). Alternatively, other variables (*e.g.* water and nitrogen availability, ambient CO_2_ concentration) and combinations thereof could result in conditions where the difference in leaf structure scales up to the whole plant level. Given the presumed hydraulic benefit of BSEs, situations where the hydraulic system is pushed to the limit (*e.g.* high evaporative demand) might be useful to maximize the benefit of BSEs. We endeavor to address these questions in the near future.

## Conclusions

The presence of BSEs in heterobaric tomato plants is coordinated with plastic variation in both structural and physiological leaf traits under different growth irradiance levels. Irradiance level altered mainly stomata pore index, minor vein density and leaf hydraulic conductance in heterobaric plants and leaf intercellular air spaces, modelled mesophyll conductance and photosynthetic assimilation rate in homobaric plants. This variation, however, allows both genotypes to maintain leaf physiological performance and growth under both irradiance conditions and results in the carbon economy and allocation of either genotype being indistinguishable within each irradiance level. Further insight into this fascinating complexity will come when the genetic basis for BSE development is unveiled.

## Acknowledgments

This work was funded by a grant (RED-00053-16) from the Foundation for Research Assistance of the Minas Gerais State (FAPEMIG, Brazil). L.E.P.P. acknowledges a grant (307040/2014-3) from the National Council for Scientific and Technological Development (CNPq, Brazil).

## Author contributions

M.A.M.B. and D.H.C. conducted experiments and prepared Figures and/or tables. A.A.A., W.L.A. D.M.R., S.C.V.M. and L.E.P.P. designed experiments, contributed reagents/materials/analysis tools and reviewed drafts of the paper. A.Z. conceived and designed the experiments, analyzed the data and wrote the paper with contributions from the other authors.

## Supplemental Information

**Fig S1**. Photosynthetic light response curves in heterobaric and homobaric cv. MT plants.

**Fig S2**. Transpiration efficiency in heterobaric and homobaric plants in the tomato cv. M82 background grown in the sun and shade.

**Fig S3**. Stomatal traits in heterobaric and homobaric cv. MT plants grown in the sun and in the shade.

**Fig S4**. Detail of mesophyll anatomy in leaves of sun-and shade-grown homobaric and heterobaric cv. MT plants.

**Fig S5**. Carbon isotope composition changes in response to irradiance in heterobaric and homobaric cv. MT plants.

**Fig S6**. Side branching ratio in MT and *obv* plants grown in the sun and shade.

**Table S1**. Photosynthetic parameters from light response curves in MT and obv grown in sun and shade

**Table S2**. Chlorophyll fluorescence analyses in MT and *obv* grown in sun and shade.

**Table S3**. Carbon isotope composition in MT and *obv* grown in sun and shade.

**Table S4**. Leaf carbohydrate and pigment content in MT and *obv* grown in sun and shade.

**Table S5**. Agronomic parameters (yield and Brix) in homobaric and heterobaric plants of tomato cultivars MT and M82 grown in the sun and shade.

**Table S6**. Gas exchange parameters determined in fully-expanded leaves of heterobaric (Micro-Tom, MT) and homobaric (*obscuravenosa*, *obv*) in two irradiance levels (sun/shade, 900/300 μmol photons m^−2^ s^−1^).

